# Poxvirus gene repertoires are shaped by expansion-biased birth-and-death dynamics and genome architectural constraints

**DOI:** 10.64898/2026.09.02.748683

**Authors:** Arnon Plianchaisuk, Spyros Lytras, Hiroyuki Hikida, Yuichiro Nakatani, Emma F. Harding, Aris Katzourakis, Jumpei Ito, Kei Sato

**Affiliations:** Laboratory of Virus Informatics, Department of Biological Informatics, Bioinformatics Center, Research Institute for Microbial Diseases, The University of Osaka, Suita, Japan; Center for Advanced Modalities and Drug Delivery Systems, The University of Osaka, Suita, Japan; Division of Systems Virology, Department of Microbiology and Immunology, The Institute of Medical Science, The University of Tokyo, Tokyo, Japan; Antigen Evolution and Design Laboratory, Department of Structural Biology and Chemistry, Institut Pasteur, Paris, France; National Institute for Infectious Diseases, Department of Virology I, Japan Institute for Health Security, Tokyo, Japan; Laboratory of Medical and Evolutionary Genomics, Department of Biological Informatics, Bioinformatics Center, Research Institute for Microbial Diseases, The University of Osaka, Suita, Japan; Department of Biology, University of Oxford, Oxford, United Kingdom; International Research Center for Infectious Diseases, The Institute of Medical Science, The University of Tokyo, Tokyo, Japan; International Vaccine Design Center, The Institute of Medical Science, The University of Tokyo, Tokyo, Japan; Graduate School of Medicine, The University of Tokyo, Tokyo, Japan; Graduate School of Frontier Sciences, The University of Tokyo, Kashiwa, Japan; Programme in Emerging Infectious Diseases, Duke-NUS Medical School, Singapore

**Author notes:** Corresponding authors (Jumpei Ito) (Kei Sato). Lead contact (Jumpei Ito).

**Keywords:** poxvirus evolution, comparative genomics, birth-and-death evolution, genome architecture, horizontal gene transfer

## Abstract

Poxviruses (family *Poxviridae*) are DNA viruses with exceptionally large genomes containing hundreds of genes. Yet the evolutionary processes contributing to their numerous gene repertoires remain insufficiently understood. Here, we inferred ortholog groups of genes across 58 poxvirus genomes and comprehensively analyzed gene gain, gene duplication, gene loss, horizontal gene transfer, and synteny conservation. Across poxviruses, coding capacity scaled linearly with genome length. Gene repertoires evolved through birth-and-death dynamics but generally expanded because gene gain cumulatively exceeded gene loss and gained genes subsequently underwent further duplication. Gene gain and duplication were concentrated in the terminal regions of most poxvirus genomes. In viral lineages whose gene repertoires expanded extensively, terminal-region expansion was accompanied by further accumulation of duplicated genes within the otherwise conserved central genomic regions. The phylogenetic placement of the bird-infecting genus *Avipoxvirus*, together with evidence that several horizontally acquired genes originated from mammals, raises the possibility of a mammalian origin for *Avipoxvirus*. Altogether, our findings reveal that poxvirus gene repertoire evolution is shaped by expansion-biased birth-and-death dynamics and genome architectural constraints.

## Introduction

Viral genomes evolve under strong constraints maximizing their coding capacity within limited genomic space. To achieve this, viruses with small genomes employ several strategies such as overlapping open reading frames and programmed ribosomal frameshifting^1^. In contrast, poxviruses stand out for their exceptionally large genomes among viruses of public health importance. Poxviruses possess double-stranded DNA genomes reaching several hundred kilobases and encoding hundreds of genes^2^. Gene repertoires of viruses with such large genomes may have evolved under constraints distinct from those acting on viruses with smaller genomes. However, the evolutionary patterns and constraints specific to poxviruses remain insufficiently understood.

Poxviruses (family *Poxviridae*) are members of nucleocytoplasmic large DNA viruses (NCLDVs), including giant virus families such as the *Mimiviridae* and *Phycodnaviridae*, as well as vertebrate pathogens such as the *Asfarviridae* and *Iridoviridae*^3^. Poxviruses infect a broad range of vertebrates and insects, and some poxviruses are causative agents of medical and veterinary diseases^4^. The most significant poxvirus diseases in humans are smallpox and Mpox, caused by the now-eradicated variola virus and the zoonotic monkeypox virus, respectively. Smallpox was one of the deadliest infectious diseases in recorded history and is estimated to have killed 300–500 million people worldwide^5^. In addition, the recent Mpox outbreak was declared a Public Health Emergency of International Concern by the World Health Organization twice in 2022 and 2024^6,7^.

The gene repertoire evolution of poxviruses and other NCLDVs has been investigated from multiple perspectives, including gene gain, gene loss, horizontal gene transfer (HGT), and genomic synteny^2,8–12^. However, these aspects have often been examined separately, limiting a comprehensive understanding of how poxvirus gene repertoires have collectively evolved over time. Furthermore, early studies were constrained by the limited availability and diversity of genome sequences^2,10^, whereas more recent studies have focused primarily on the genus *Orthopoxvirus*^11,12^, which includes variola virus and monkeypox virus. Altogether, these limitations have left family-wide and lineage-specific patterns of poxvirus gene repertoire evolution insufficiently understood.

Here, we inferred orthologous relationships among genes in 58 poxvirus genomes representing diverse viral lineages and host taxa, covering the entire *Poxviridae* family. On this basis, we reconstructed gene gain, duplication, loss, and HGT across the poxvirus phylogeny. Our study provides novel insights into the mechanisms underlying gene repertoire and genome evolution in poxviruses, with broader implications for large DNA viruses.

## Results

### Poxvirus ortholog groups and phylogeny

To reconstruct the evolutionary history of the poxvirus gene repertoire, genomes of 58 poxviruses—including those of four monkeypox viruses representing four distinct subclades—were retrieved from the NCBI RefSeq^13^ and GenBank^14^ databases (**Supplementary Table 1**). We extracted pre-annotated genes from these genomes and clustered them into ortholog groups using OrthoFinder2^15,16^. Ortholog groups are sets of homologous genes, including both orthologs and paralogs (i.e., duplicated genes), that descended from a single ancestral gene^15^. To recover genes missing from pre-defined gene annotations, we performed tblastn searches against poxvirus genomes to identify open reading frames homologous to known poxviral genes. Because poxvirus genes lack introns, this approach allows accurate recovery of full-length gene sequences. Newly identified genes were assigned to the same ortholog group as their top-hit gene for enabling more complete orthology inference (**see Methods**). In total, we identified 1,553 ortholog groups comprising 11,699 homologs across the 58 analyzed genomes. The number of ortholog groups per genome ranged from 113 to 269, and the total number of genes per genome ranged from 120 to 334 (**Figure 1A; Supplementary Figure 1A**).

**Figure 1.**
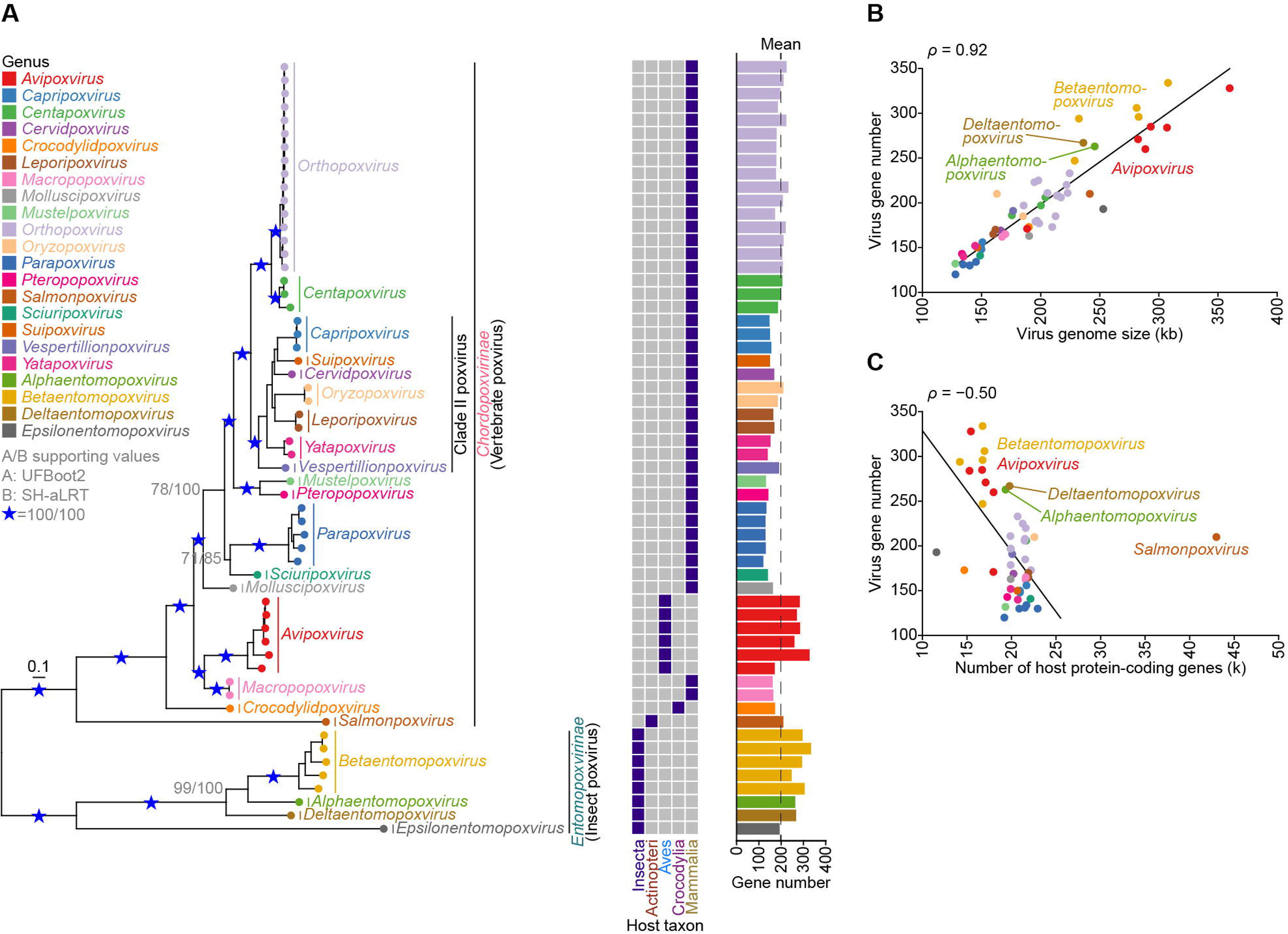
Variation in gene number and genome size among poxviruses. (**A**) Phylogenetic tree of poxvirus lineage (left), taxonomic class of the documented hosts (middle), and number of genes in each poxvirus (right). Nodes with both ultrafast bootstrap (UFBoot2) and Shimodaira–Hasegawa-like approximate likelihood ratio test (SH-aLRT) support values of 100 are indicated by blue stars, while all other nodes are labeled with their support values in grey. (**B and C**) Scatter plots showing the relationship between the number of poxvirus genes and (**B**) poxvirus genome size (kilobase; kb) or (**C**) the number of host protein-coding genes. The fitted linear regression line and Spearman’s rank correlation coefficient (ρ) are shown. For panel **C**, poxviruses associated with multiple host species were assigned the average number of protein-coding genes across their documented hosts. Salmon gill poxvirus was excluded from the correlation and linear regression analyses because its host, Atlantic salmon (*Salmo salar*), has an unusually large number of protein-coding genes resulting from a lineage-specific whole-genome duplication event^64^.

To infer the evolutionary relationships among the 58 analyzed poxviruses, a viral species phylogeny was reconstructed based on 22 single-copy ortholog groups conserved across all examined genomes (**Figure 1A; Supplementary Table 2**). The reconstructed tree was well supported, with 82.1% and 89.3% of nodes receiving support values > 95 in the ultrafast bootstrap approximation^17^ and > 80 in the Shimodaira-Hasegawa-like approximate likelihood ratio test^18^, respectively. Consistent with previous studies^2,10,19^, poxviruses were divided into the *Chordopoxvirinae* and *Entomopoxvirinae* subfamilies, whose members are known to infect vertebrates and insects, respectively. The phylogeny of viruses within *Chordopoxvirinae* broadly reflected the phylogeny of their hosts, except for the avian-infecting *Avipoxvirus* genus, which formed a clade with the marsupial-infecting *Macropopoxvirus* genus. *Orthopoxvirus*, including variola virus and monkeypox virus, was most closely related to *Centapoxvirus*, followed by a clade comprising Clade II poxviruses^10^. We simply refer to all viruses within the latter clade as Clade II poxviruses.

We next examined poxvirus gene number, genome size, and their correlation. The number of poxvirus genes varied across genera (**Figure 1A**). Avipoxviruses and entomopoxviruses have substantially more genes than poxviruses from other genera. Viral gene number showed a strong positive correlation with genome size (Spearman’s rank correlation coefficient = 0.92), where poxviruses with larger genomes, such as avipoxviruses and entomopoxviruses, encoded more genes (**Figure 1B**). This correlation remained statistically significant after accounting for phylogenetic dependence (**Supplementary Figure 1B**). These results indicate that the coding capacity of poxviruses increases linearly with genome length.

In addition, poxvirus gene number was negatively correlated with host protein-coding gene number and host genome size (**Figure 1C; Supplementary Figures 1C and 1D**). Notably, hosts of avipoxviruses and entomopoxviruses—birds and insects, respectively—generally have relatively smaller genome size compared to mammals^20,21^. These findings collectively suggest a potential evolutionary association among virus and genomic features of their hosts.

### Ortholog group gain and loss events and their evolutionary dynamics

To reconstruct the evolutionary trajectories of gene repertoires in poxviruses, we inferred ortholog group gain and loss events along branches of the poxvirus phylogenetic tree using three ancestral state reconstruction methods (**Figure 2A; Supplementary Figures 2A and 2B**). Here, gain and loss events refer to the acquisition or loss of an entire ortholog group, rather than changes in gene copy number within an ortholog group. Only transitions consistently inferred by all three methods were considered as gain and loss events.

**Figure 2.**
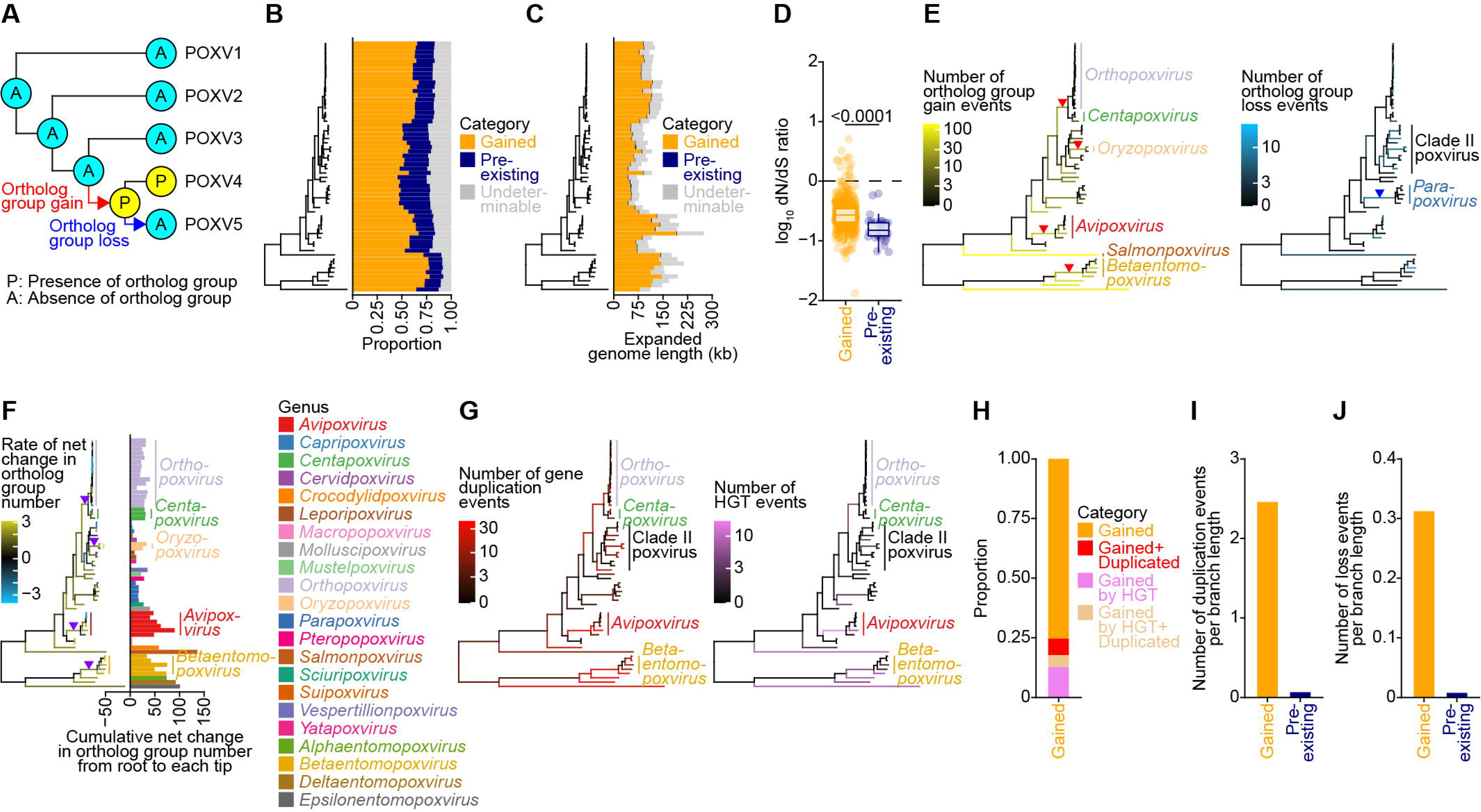
Gene gain, duplication, and loss during poxvirus evolution. (A) Inferring ortholog group gain and loss events. Ortholog group gain and loss events were inferred by reconstruction of the presence (P) or absence (A) of each ortholog group at internal nodes and subsequent identification of state transitions along branches. (B) Proportions of gained, pre-existing, and undeterminable ortholog groups in each poxvirus genome. (C) Contributions of gained and pre-existing ortholog groups to poxvirus genome expansion. Expanded genome length attributable to gene gain and gene duplication are shown. Only non-overlapping genetic regions were considered in the calculation. (D) Distribution of per-ortholog-group dN/dS ratios of gained and pre-existing ortholog groups. (E) Numbers of ortholog group gain (left) and loss (right) events in each phylogenetic branch. Branch colors indicate event counts on a pseudo-logarithmic scale. Red and blue triangles indicate stem lineages of interest. (F) Rates of net change in ortholog group number in each phylogenetic branch (left) and cumulative net changes from the MRCA to each tip (right). Branch colors indicate transformed rates, calculated as sign(x) × log_10_(|x| + 1), where x denotes the original rate. Violet triangles indicate stem lineages of interest. (G) Numbers of gene duplication (left) and HGT (right) events in each phylogenetic branch. (H) Proportions of gained ortholog groups expanding poxvirus genomes stratified by whether they further underwent gene duplication and whether they were HGT-derived. (**I and J**) Numbers of duplication (**I**) and loss events (**J**) per branch length in gained and pre-existing ortholog groups. For each category, the total event counts were normalized by the total length of branches descending from MRCAs of the corresponding ortholog groups. Only branches on which the ortholog groups were continuously present were included.

We identified 1,066 gain and 204 loss events of ortholog groups that occurred during poxvirus evolution. Across all analyzed poxviruses, an average of 60.2% of ortholog groups were gained after the most recent common ancestor (MRCA) of poxviruses emerged (**Figure 2B**). Furthermore, gained ortholog groups accounted for 70.2% of the expanded genome length of all poxviruses, suggesting that gene gain has been the primary driver of gene repertoire expansion and, consequently, that of genome expansion in the poxvirus evolution (**Figure 2C**). Although dN/dS ratios were generally less than 1, they were higher in gained ortholog groups than in pre-existing ones (**Figure 2D**). Moreover, a subset of gained ortholog groups exhibited dN/dS ratios greater than 1 (**Figure 2D**). These results suggest that gained ortholog groups are generally subject to more relaxed evolutionary constraints, with some potentially experiencing diversifying selection.

Next, we counted gain and loss events of ortholog groups along each branch of the poxvirus species tree (**Figure 2E; Supplementary Figures 2A and 2B**). The number of gain events was relatively higher in the stem lineages of *Avipoxvirus*, *Betaentomopoxvirus*, *Oryzopoxvirus*, and the clade comprising *Orthopoxvirus* and *Centapoxvirus*, as well as in the long branches leading to *Salmonpoxvirus* and other entomopoxviruses (**Figure 2E, left; Supplementary Figure 2A**). On the other hand, the number of loss events was relatively higher in the stem lineage of *Parapoxvirus* and some sublineages within Clade II poxviruses (**Figure 2E, right; Supplementary Figure 2B**).

To reconstruct the evolutionary dynamics of ortholog group number, we calculated the net change in ortholog group number for each phylogenetic branch, defined as the number of gain events minus the number of loss events. We also calculated the rate of this net change by normalizing the change by the branch length. Although the rate varied across the phylogenetic tree, the cumulative net change from the root to each tip was positive across all analyzed poxviruses (**Figure 2F; Supplementary Figure 2C**). This result suggests that the number of ortholog groups has cumulatively increased across all extant poxvirus lineages. Particularly, the elevated rate of the net change was observed along the branches with higher number of gain events described earlier, suggesting that gain events outpaced loss events along these branches and thereby drove gene repertoire expansion (**Figure 2F**).

### Gene duplication, horizontal gene transfer, and gene loss events in poxviruses

To elucidate the mechanisms underlying poxvirus gene repertoire expansion, we inferred gene duplication and horizontal gene transfer (HGT) events that occurred during poxvirus evolution (**Figure 2G**). Gene duplication events were identified using the duplication-loss-coalescence analysis implemented in OrthoFinder2^16^. HGT events were inferred by comparing poxvirus proteins from gained ortholog groups with non-poxvirus proteins available in the RefSeq database (**see Methods**). Among the gained ortholog groups, 17.7% were inferred to be HGT-derived, whereas 12.3% further underwent gene duplication (**Figure 2H**). Most duplication and loss events involved gained ortholog groups rather than pre-existing ones, indicating a sequential pattern of gene gain followed by gene duplication or gene loss, consistent with the birth-and-death model of gene repertoire evolution (**Figures 2I and 2J**).

### Genomic distribution of gained ortholog groups

Next, we investigated the genomic locations of poxvirus ortholog groups. Consistent with a previous report^2^, ortholog groups in the central genomic regions of mammalian poxviruses, such as *Orthopoxvirus*, *Centapoxvirus*, Clade II poxviruses, and *Parapoxvirus*, are highly conserved while maintaining overall syntenic organization (**Figure 3A**). We therefore defined central and terminal regions of each poxvirus genome (**Supplementary Figure 3A**) and investigated genomic distribution of gained and pre-existing ortholog groups in these regions (**see Methods**). Our results further showed that gained ortholog groups accumulated in terminal regions more than in central regions in most viruses (**Figures 3B and 3C**). Although both ends of poxvirus genomes contain inverted terminal repeats (ITRs)^22^, the terminal regions associated with gene repertoire expansion extended beyond the boundaries of these ITRs (**Figure 3D; Supplementary Figure 3B**). In addition, genes belonging to some gained ortholog groups have undergone repeated local duplication within individual terminal regions, further contributing to gene repertoire expansion in terminal regions (**Figure 3D; Supplementary Figure 3B**).

**Figure 3.**
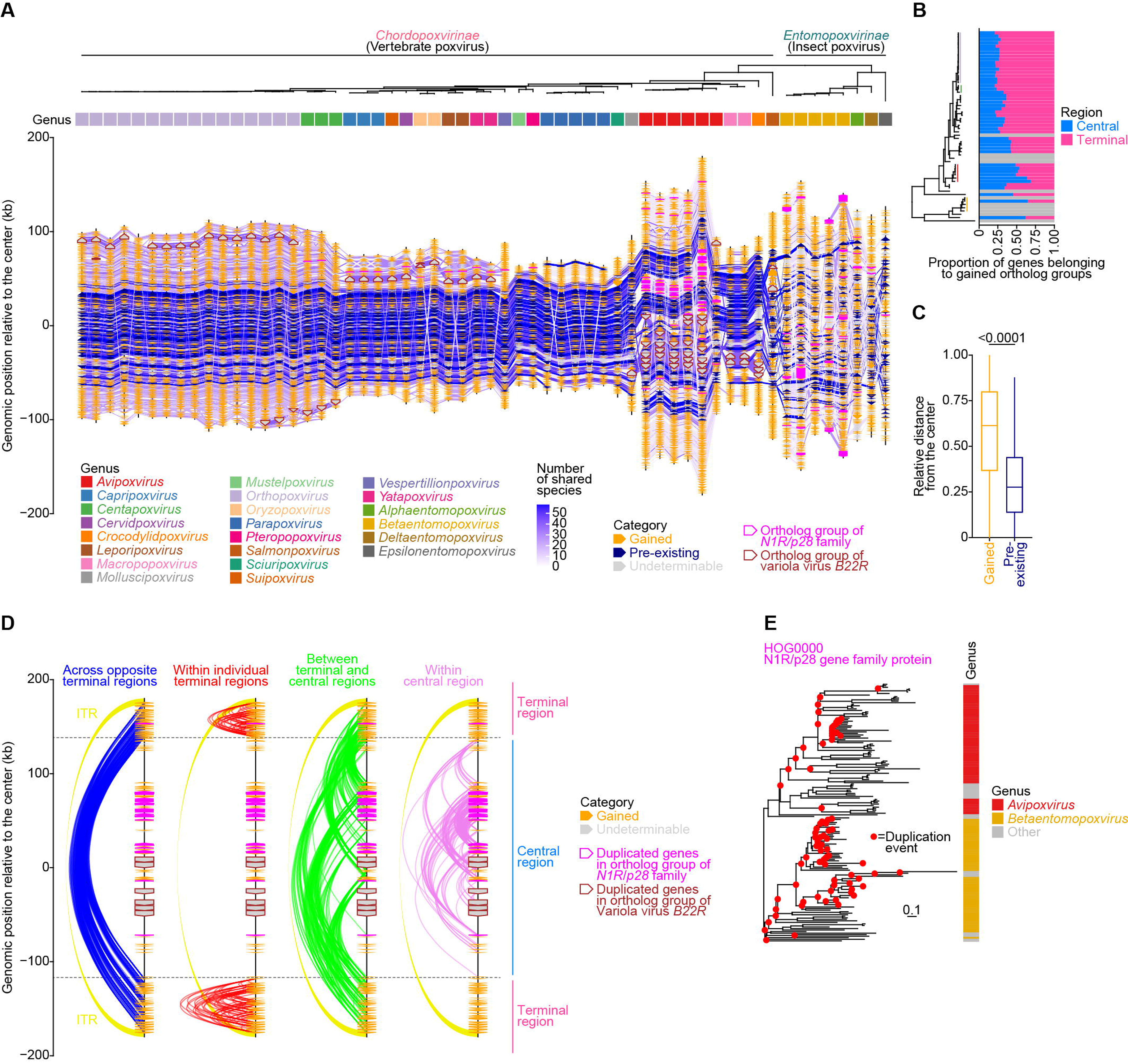
Genomic location of poxvirus ortholog groups. (A) Genomic synteny of poxvirus genomes. Orange, navy blue, and light grey arrows indicate gained, pre-existing, and undeterminable ortholog groups, respectively. Ribbons connecting ortholog groups indicate orthologous relationships, and ribbon colors represent the number of poxvirus genomes sharing each ortholog group. Genomic positions are shown relative to the genome center, which was set to 0. Ortholog groups of *N1R/p28* family and variola virus *B22R* are outlined in brown and magenta, respectively (B) Proportions of genes belonging to gained ortholog groups in central and terminal genomic regions. Boundaries between central and terminal regions were defined using a sliding-window approach based on the proportion of genes belonging to pre-existing ortholog groups. Rows in grey indicate that central and terminal genomic regions cannot be defined by the sliding window method. (C) Relative distance of gained and pre-existing ortholog groups from the center of poxvirus genomes. (D) Duplicated genes in all ortholog groups in the genome of canarypox virus. Genomic positions are shown relative to the genome center, which was set to 0. Line colors indicate genomic distribution of paralogous relationships between duplicated genes. Duplicated genes in ortholog groups of *N1R*/*p28* family and variola virus *B22R* are outlined in brown and magenta, respectively. ITRs are shown by yellow ribbons. (E) Gene tree of an ortholog group of *N1R/p28* family. Nodes in red indicate gene duplication events.

Beyond the expansion of terminal regions described above, duplicated genes also accumulated extensively within the central genomic regions of avipoxviruses and betaentomopoxviruses, whose gene repertoires and genomes expanded extensively (**Figure 3D; Supplementary Figure 3B**). Duplicated genes in central genomic regions belong to a few ortholog groups. A prominent example was the extensive duplication of the *N1R/p28* family, the members of which encode ubiquitin ligases targeting substrates in poxvirus viral factories^23,24^ (**Figure 3D; Supplementary Figure 3B**). This family expanded convergently within the central genomic regions of avipoxviruses and betaentomopoxviruses (**Figure 3E; Supplementary Figure 3B**). Similarly, variola virus *B22R* family, which functions in T cell inactivation^25^, have undergone substantial expansion within the central genomic regions of avipoxviruses (**Figure 3D; Supplementary Figure 3B**). Notably, duplicated genes in central genomic regions accounted for 16.6% and 15.8% of the total genome length in avipoxviruses and a betaentomopoxvirus, respectively. Altogether, these results suggest that gene repertoires and genomes of these poxvirus lineages have expanded additionally through accumulation of duplicated genes within central genomic regions.

### Putative donors of poxvirus HGT-derived genes and association with hosts

Although previous studies have investigated HGT events in poxviruses, the taxonomic relationship between gene donors and known poxvirus hosts remains unclear^9,10,26,27^. Therefore, we systematically examined the taxa of putative donors of 159 HGT-derived ortholog groups using a BLAST-based inference method (**Figure 4A; Supplementary Figure 4; Supplementary Table 3; see Methods**). The closest matches for HGT-derived ortholog groups in mammalian chordopoxviruses were predominantly mammalian genes. Likewise, those in entomopoxviruses, which infect insects, were predominantly insect genes. These findings support an evolutionary scenario in which poxvirus genes were gained horizontally from their extant or ancestral hosts rather than from unrelated organisms. We further examined the proportion of HGT-derived ortholog groups for which the taxa of the putative donors matched those of the known host at different taxonomic ranks. The degree of concordance increased progressively toward higher taxonomic ranks, from species to phylum (**Figure 4B**). This pattern is consistent with two non-mutually exclusive scenarios. First, HGT occurred during long-term coevolution with ancestral host lineages rather than during recent interactions with extant hosts. Second, subsequent divergence of the transferred genes in viruses and their homologs in host lineages may have reduced the taxonomic resolution of putative donor inference.

**Figure 4.**
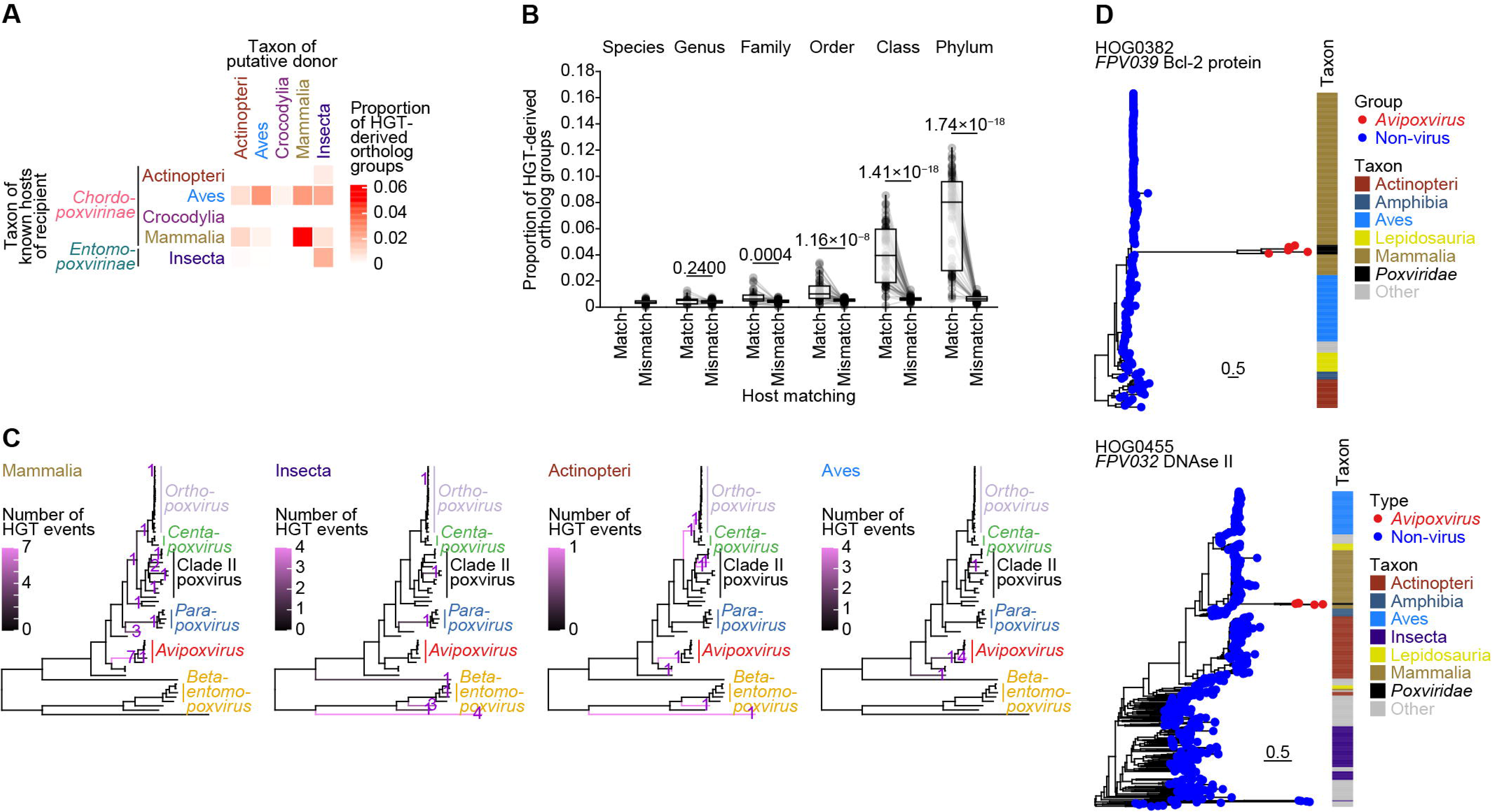
Putative donors of horizontally transferred genes in poxviruses. (A) Host–donor taxonomic associations of genes horizontally transferred to poxviruses. For each taxon of hosts of recipient poxviruses (row), we calculated the proportion of HGT-derived ortholog groups inferred to have originated from putative donors belonging to each taxon (column). For each donor–recipient pair, the proportion was calculated by dividing the sum of the numbers of HGT-derived ortholog groups in recipient poxviruses by the sum of the numbers of ortholog groups in those viruses. Heatmap colors represent the calculated proportions. (B) Proportions of HGT-derived ortholog groups stratified by whether the putative donor taxon matched the documented host taxon at different taxonomic ranks. (C) Numbers of HGT events along branches of the poxvirus species tree, stratified by putative donor class. (D) HGT-derived ortholog groups gained in the stem lineage of *Avipoxvirus*, including *FPV382* (upper) and *FPV032* (lower). Fowlpox virus gene names are shown for each ortholog group. Gene trees were rooted using genes from fish in class Actinopteri and those from a clade comprising protists in phyla Discosea, Evosea, and Preaxostyla, respectively.

Next, we inferred the branches of the poxvirus species tree on which HGT-derived ortholog groups were acquired (**Figure 4C**). Most mammal-derived ortholog groups found in mammalian poxviruses were inferred to have been acquired during the diversification of mammalian poxviruses rather than along the ancestral poxvirus lineage. A similar pattern was observed for insect-derived ortholog groups in insect poxviruses.

Notably, the BLAST-based inference suggested that the *Avipoxvirus* genus harbors comparable proportions of mammalian-derived and avian-derived ortholog groups (**Figure 4A**). This pattern was further supported by the analysis of HGT branches, which showed that eight mammalian-derived and five avian-derived ortholog groups were acquired during the emergence and early diversification of the *Avipoxvirus* genus (**Figure 4C**). To further examine this, we reconstructed a gene tree for each of the eight ortholog groups inferred to be of mammalian origin, together with its homologs from a diverse range of organisms (**Figure 4D**). This analysis further supported a mammalian origin for two of these ortholog groups, related to fowlpox virus Bcl-2 and DNase II proteins. Together with the phylogenetic placement of *Avipoxvirus* within a clade predominantly composed of mammalian poxviruses (**Figure 1A**), these results raise the possibility that the stem lineage of *Avipoxvirus* was associated with mammalian hosts (**see Discussion**).

### Protein domain gain and loss events and functional enrichment

To characterize changes in functional genomic content across poxvirus evolution, we reconstructed gain and loss events at the protein domain level (**Figure 5A; Supplementary Figure 5**). We then performed an enrichment analysis to identify domains overrepresented among gained and lost ortholog groups (**Figure 5B; Supplementary Tables 4 and 5**). The poxvirus Bcl-2-like domain was significantly enriched among these ortholog groups (adjusted *P* = 0.0004, odds ratio = 3.21) (**Supplementary Table 5**). Gene gain and loss events involving this domain occurred across 11 ortholog groups in the clade comprising *Orthopoxvirus*–*Centapoxvirus* and Clade II poxviruses (**Figures 5C**). Genes belonging to these gained ortholog groups encode proteins consisting solely of the poxvirus Bcl-2-like domain and collectively constitute the poxvirus Bcl-2-like gene family. Some members of this family are known to have immunomodulatory functions^28–31^ (**Figure 5C**). All genes in this family were located in terminal genomic regions, as described in a previous study^32^ (**Figure 5D**). Genes in this family also tended to exhibit higher dN/dS ratios than pre-existing ortholog groups (**Figure 5E**). Moreover, duplication and loss events occurred independently in five and six of the 11 ortholog groups, respectively. Together, these findings suggest that the poxvirus Bcl-2-like gene family has undergone birth-and-death evolution, characterized by gene gain followed by recurrent duplication or loss under relatively relaxed selective constraint.

**Figure 5:**
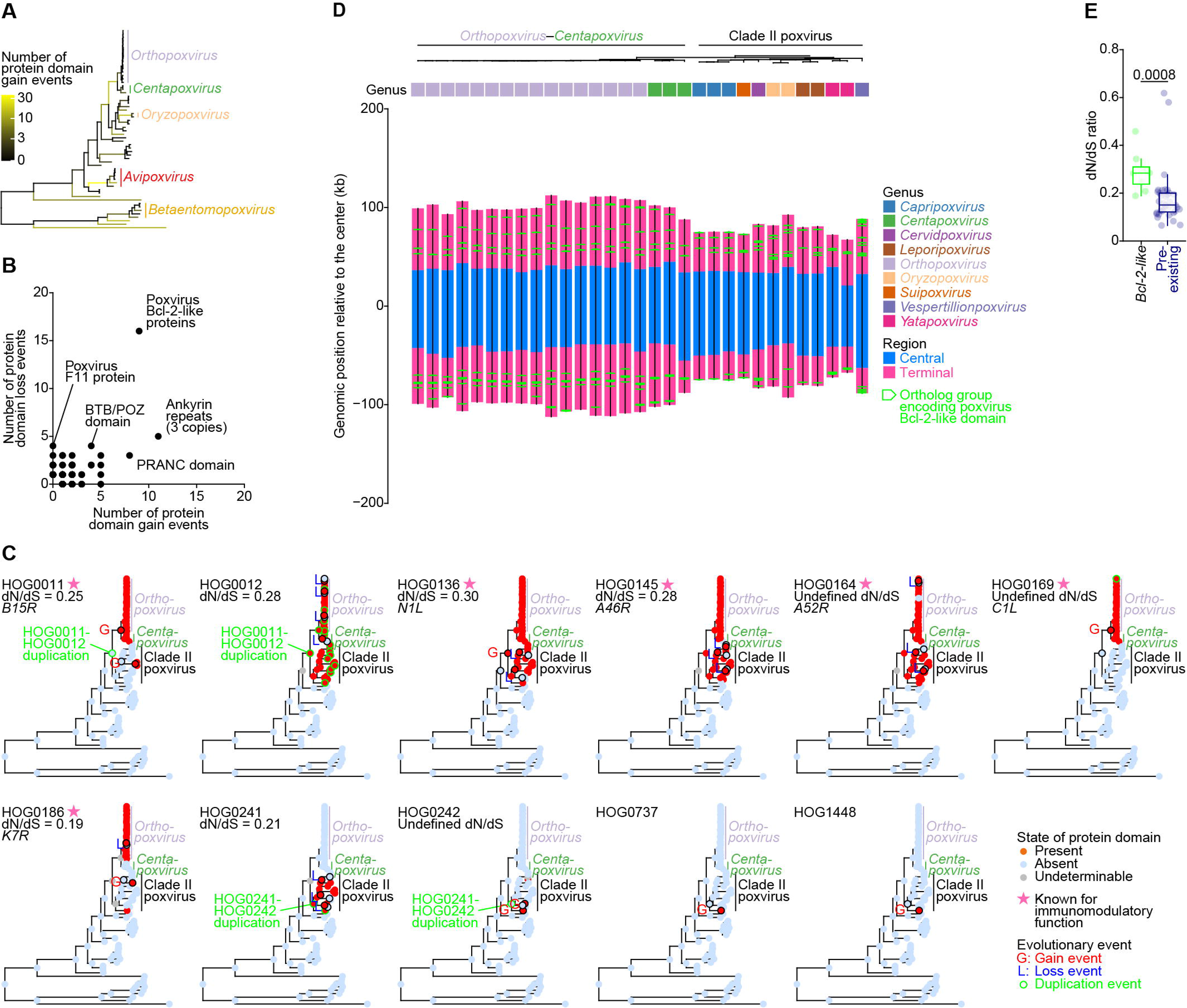
Protein domains gained during poxvirus evolution. (A) Number of protein domain gain events inferred for each branch of the poxvirus species tree. Only those accompanied by ortholog group gain events were counted. Branch colors represent event counts on a pseudo-logarithmic scale. (B) Numbers of gain and loss events associated with each protein domain. The top three protein domains with the highest number of gain or loss events are labelled. (C) Gain, duplication, and loss events of ortholog groups whose gene members encode poxvirus Bcl-2-like protein domains. G: gain event; L: loss event. Nodes where gene duplication occurred are outlined in yellow. Pink stars indicate ortholog groups containing vaccinia virus genes known for their immunomodulatory function. (D) Genomic location of ortholog groups encoding poxvirus Bcl-2-like protein domains, outlined in green. (E) Distribution of dN/dS ratios of ortholog groups whose gene members encode Bcl-2-like protein domains and pre-existing ortholog groups.

## Discussion

In this study, we performed orthology-based comparative genomic analyses of 58 poxvirus genomes representing diverse viral lineages and host taxa to investigate gene repertoire evolution across the poxvirus family. We found that the coding capacity of poxviruses increases linearly with genome length (**Figure 1**). We also revealed a long-term trend toward gene repertoire expansion during poxvirus evolution (**Figure 2**). This expansion exhibited two spatial patterns, including terminal-region expansion broadly observed across the family and additional central-region expansion in specific lineages (**Figure 3**). Our results further suggest that some HGT events occurred during long-term associations with ancestral host lineages (**Figure 4**). Altogether, by connecting various aspects of genome evolution that have often been examined separately^2,8–12^, our study provides an integrated insights into family-wide and lineage-specific patterns of poxvirus gene repertoire evolution.

We showed that gained ortholog groups have undergone duplication and loss more frequently than pre-existing ones (**Figures 2I and 2J**) and that their constituent genes evolved under relatively relaxed selective constraints (**Figure 2D**). This evolutionary pattern was exemplified by recurrent duplication and subsequent loss of Bcl-2-like genes (**Figure 5C**). Such a pattern is comparable to the birth-and-death evolution of gene families widely observed in cellular organisms^33^. Importantly, the birth-and-death cycle in poxviruses has been biased toward expansion. Ortholog group gains cumulatively outnumbered losses across the poxvirus phylogeny, and the majority of ortholog groups present in extant genomes were acquired after the poxvirus MRCA emerged. In a previous study, the genomic accordion model was proposed based on experimental evolution to explain how poxviruses rapidly adapt to strong selective pressures^34^. The model posits that transient gene amplification and diversification are followed by the loss of excess copies and the retention of one advantageous copy, resulting in no net gene repertoire expansion. Viewed over macroevolutionary timescales, our findings raise the possibility that the accordion-like cycles were not fully reversible, cumulatively resulting in a net accumulation of genes in poxvirus genomes. Altogether, these results characterize poxvirus gene repertoire evolution as an expansion-biased form of birth-and-death evolution. This evolutionary mode may also contribute to the expansion of gene repertoires documented in several other NCLDV groups^3^.

Our analyses revealed two spatial patterns of gene repertoire expansion in poxviruses (**Figure 3**). The first pattern involves accumulation of gained ortholog groups in terminal regions and is characterized by both ITR-mediated gene duplication and local gene duplications extending beyond the ITR boundaries. This pattern is common across poxviruses and consistent with the patterns documented in some linear NCLDVs^2,35^. The second pattern involves accumulation of duplicated genes in the highly conserved central genomic regions. This pattern was additionally observed in the *Avipoxvirus* and *Betaentomopoxvirus* genera, the gene repertoires of which expanded extensively. Most of these central gene clusters arose through extensive duplication of genes belonging to a small number of ortholog groups, such as the *N1R/p28* family and variola virus *B22R* family. Spatially localized gene-family expansion is common in cellular genomes and has been linked to features of genome architecture^36^. Therefore, elucidating how viral genome architecture, genome packaging, and gene regulation permit such spatial expansion would be important for understanding the mechanisms shaping poxvirus genome evolution.

Although the poxvirus species tree largely reflected a virus–host co-speciation pattern, *Avipoxvirus*, a genus of avian poxviruses, represented a striking exception. *Avipoxvirus* was nested within a clade of mammalian poxviruses (**Figure 1A**). Furthermore, our analysis of the putative donors of HGT-derived ortholog groups showed that *Avipoxvirus* genomes contained comparable numbers of mammalian-derived and avian-derived ortholog groups (**Figure 4**). Together, these findings suggest that the *Avipoxvirus* lineage may have originated from a mammal-associated ancestor and subsequently shifted its host range to avian hosts. This hypothesis is supported by a previous study demonstrating replicability of an avipoxvirus in a hamster-derived cell line^37^. Intriguingly, the *Avipoxvirus* genus has also undergone one of the most extensive gene repertoire expansions among all poxvirus lineages. Therefore, the inferred host shift may have imposed new selective pressures that contributed to this remarkable expansion. In addition, the relatively smaller number of protein-coding genes in avian genomes—and the possible absence of some host genes involved in poxvirus replication—may have contributed to the expansion of *Avipoxvirus* gene repertoires.

Several limitations of this study should be acknowledged. First, incomplete lineage sorting^38^ and genetic recombination^39^ might cause incorrect inference of the placement and timing of ancient gain, loss, and duplication events. Likewise, it can be sometimes difficult to distinguish the acquisition of a novel ortholog group from the duplication of an existing gene. For example, OrthoFinder divided the poxvirus Bcl-2-like gene family into multiple ortholog groups, although these groups are likely to descend from a common ancestral gene (**Figure 5C**). Second, our HGT inference relied on homology searches against extant proteins, which may not identify the true ancestral donor lineages accurately. For instance, several ortholog groups in chordopoxviruses inferred to be insect-derived may represent false-positive HGT inferences. Finally, although our results support a mammalian-associated ancestry of *Avipoxvirus*, this evidence remains indirect, and the hypothesis therefore requires further validation.

Despite these limitations, this study extensively provides an integrated analysis of gene repertoire evolution in poxviruses, combining evidence from dynamics of gene gain, gene duplication, gene loss, HGT involving hosts, and synteny conservation. Our findings reveal the distinctive evolutionary dynamics of poxvirus genomes and provide new insights into how *Poxviridae*, including several highly pathogenic viruses, has adapted to its hosts.

## Methods

### Data collection

Genomic data from 58 poxviruses, including those from four monkeypox viruses were retrieved from the NCBI GenBank and RefSeq databases^13,14^ on 8 October 2024 (**Supplementary Table 1**). Data on poxvirus–host relationships were retrieved from the GenomeNet Virus-Host database^40^ on 27 March 2025 (**Supplementary Table 6**). Genomic data of hosts or animals phylogenetically closely related to the hosts available in public databases were retrieved from the NCBI GenBank, NCBI RefSeq, and DNA Zoo^41^ databases on 23 February 2026 (**Supplementary Table 6**).

### Ortholog group inference of poxvirus genes

Phylogenetically hierarchical ortholog groups were inferred from the orthologous relationship among the collected poxvirus proteins using OrthoFinder v2.5.5^15,16^. The option -y was used to split paralogous clades within nested hierarchical ortholog groups into separate hierarchical ortholog groups. Only the longest protein isoform of each ortholog group was used in the orthology inference. Each of the proteins unassigned to any ortholog groups, except for those identified as phylogenetically misplaced by OrthoFinder, was designated as an ortholog group with a single protein member. Ortholog groups were named based on gene names of vaccinia virus.

To identify potential orthologs missed in existing poxvirus genome annotations, we searched poxvirus genomes using tblastn, implemented in NCBI standalone BLAST+ v2.15.0^42^, and protein sequences from each ortholog group as queries. An alignment hit was considered a putative ortholog if it had the highest bit score and met the percentage identity > 50%, query coverage per high-scoring segment pair > 95%, and *E*-value ≤ 1 10^−5^ criteria. Translated coding sequences of putative orthologs with start and stop codons, whose lengths are at least 30 amino acids, were then predicted using EMBOSS getorf v6.6.0.0^43^. An ortholog was considered present if (1) the best-matching ortholog group for the longest translated coding sequences corresponding to the putative ortholog matched the ortholog group whose assigned proteins were used in the tblastn search to identify the gene and (2) the protein corresponding to the putative ortholog had alignment hits with at least 50% of the proteins assigned to the best-matching ortholog group. The translated coding sequences were aligned to assigned proteins using Diamond v2.1.8^44^.

### Reconstruction of the phylogenetic tree of poxvirus lineage

From the 1,553 identified ortholog groups, we selected 22 single-copy ortholog groups conserved in the 58 poxvirus genomes to reconstruct a phylogenetic tree (**Supplementary Table 2**). First, proteins from each of the selected ortholog groups were subject to multiple sequence alignment using MAFFT v7.505^45^ with the E-INS-I method and 1,000 maximum iterative refinements. Gaps were automatically removed using Trimal v1.4.rev22^46^. Then, the trimmed alignments were concatenated into a supermatrix containing 11,091 amino acid sites, including 9,510 parsimony-informative sites. Finally, the phylogenetic tree was reconstructed from the supermatrix using the maximum-likelihood-based IQ-TREE v2.3.4^47^. The best-fit amino acid substitution models and partitioning schemes were selected automatically using ModelFinder^48^ and PartitionFinder^49^ implemented in the IQ-TREE suite, respectively. Branch support was assessed using ultrafast bootstrap approximation^17^ and Shimodaira-Hasegawa-like approximate likelihood ratio test^18^ with 1,000 bootstrap replicates.

### Gene gain and loss analysis

Ortholog group gain and loss events were estimated by reconstructing the ancestral state of presence or absence of orthologs for each ortholog group across poxvirus ancestors. First, the state for the presence (P) or absence (A) of each ortholog group was assigned to each of the 58 poxviruses. Then, the states for the presence or absence of that ortholog group across poxvirus ancestors were estimated using three ancestral state reconstruction methods, comprising the marginal posterior probability approximation (MPPA), maximum a posteriori (MAP), and maximum-parsimony-based DOWNPASS methods, all of which are implemented in PastML v.1.9.50^50^. The gain and loss events of ortholog groups were then inferred from the state changes A to P and P to A from an ancestral node to its immediate descendant node, respectively. We then counted ortholog group gain and loss events at each branch of the phylogenetic tree. To calculate the rate of net ortholog group gain at each branch, we calculated the net change in ortholog group number by subtracting the number of ortholog group loss events from that of ortholog group gain events and subsequently divided the net change by the branch length.

### Gene origin analysis

Ortholog groups were categorized into gained, pre-existing, and undeterminable categories based on the presence and absence of an ancestral orthologous gene in the MRCA of poxviruses. The gained category was assigned if the ortholog group was absent in the MRCA of poxviruses and present in at least one poxvirus genome analyzed. The pre-existing category was assigned if the ortholog group was present in the MRCA of poxviruses analyzed. The undeterminable category was assigned if the presence or absence of the ortholog group in the MRCA of poxviruses analyzed was undeterminable.

Gene duplication events were identified using the duplication-loss-coalescence analysis implemented in OrthoFinder with default parameters^16^.

To inferred HGT events, first, ortholog groups derived from gene duplication were pooled to reduce the influence of gene duplication in downstream analyses. The pooled ortholog groups were categorized into gained, pre-existing, and undeterminable ortholog groups using the methods explained above. Only gained ones were analyzed for HGT events that occurred after the emergence of the MRCA of poxviruses. The putative donor for each of the gained pooled ortholog groups was inferred by comparing amino acid sequences of poxvirus proteins from the pooled ortholog group against those of all curated proteins deposited in the RefSeq database release 229, using BLASTp implemented in BLAST+ v2.15.0. Each pooled ortholog group was considered derived by HGT if at least half of the protein members could be aligned to the non-poxvirus proteins and satisfied the criteria of a query coverage per high-scoring segment pair > 50% and an *E*-value ≤ 1 10^−5^ (**Supplementary Figure 6**). Only the best alignment hits, defined as those with the highest bit scores, were selected for identifying the putative donors. The proportions of HGT-derived ortholog groups (**Figures 4A and 4B; Supplementary Figure 4**) were calculated relative to the total number of HGT-derived pooled ortholog groups.

Next, we inferred the representative taxonomic class of putative donors for each pooled ortholog group by using a majority-voting approach. A class was assigned when at least half of the proteins in the pooled ortholog group were inferred to have putative donors from the same class. This approach could identify the taxonomic class of putative donors of 126 out of 159 HGT-derived pooled ortholog groups (**Supplementary Table 3**).

### Gene tree analysis of HGT-derived genes

Gene trees were then reconstructed to provide the phylogenetic evidence of HGT. First, for each ortholog group, the best-hit proteins for each protein from each ortholog of poxvirus ortholog groups were retrieved. Then, homologs of the best-hit proteins were searched against all RefSeq proteins using BLASTp. The homologs that met the criteria of a query coverage per high-scoring segment pair > 90% and an *E*-value ≤ 1 10^−5^ were selected for the gene tree reconstruction. Gene trees were reconstructed using the same methods used for the reconstruction of the poxvirus phylogenetic tree, except for the partitioning scheme that was not applied. Gene trees were rooted manually according to the current systematic knowledge.

### Selective pressure analysis

The nucleotide and amino acid coding sequences of homologs in 626 poxvirus ortholog groups, each of which contains at least three homologs, were subject to multiple sequence alignment using MAFFT with the E-INS-I method and 1,000 maximum iterative refinements. Codon alignments were then created from the nucleotide and amino acid alignments using PAL2NAL v14^51^. Selective pressure across poxvirus homologs in each ortholog group was then estimated using the dN/dS ratio (i.e., the ratio between the number of non-synonymous substitutions per non-synonymous site to the number of synonymous substitutions per synonymous site) calculated by the kaks function implemented in the seqinr R package v4.2.36^52^.

### Gene synteny analysis

To compare genomic positions of ortholog groups across poxvirus genomes, genome coordinates were centered by setting the midpoint of each genome to zero. For each ortholog group, the relative distance to the genomic center was calculated as the mean distance between the center of each homologous member to the genomic center.

Assuming that central regions of poxvirus genomes are enriched in pre-existing ortholog groups, we identified boundaries between central and terminal regions using a sliding window approach. First, we defined the middle 20% of each genome as the central reference region. We then divided each genome into overlapping windows, which cover 10% of its genome length and moved in 1% increments. For each window, we calculated the proportion of homologs of pre-existing ortholog groups whose center fell within the window. The proportion was calculated only for windows containing at least ten homologs belonging to gained or pre-existing ortholog groups. The proportion in each window, whose midpoint at ≤ 35% or ≥ 65% of genome length, was compared with that in the central reference region using one-sided Fisher’s exact test. *P* values were adjusted using the Benjamini–Hochberg method, with a significance threshold of 0.05. Boundaries were inferred when at least five consecutive windows showed significant decrease in the proportion. The left and right boundaries were placed immediately inward of the midpoint of the innermost significant window on each side.

Next, we identified ITRs in each poxvirus genome. First, each genome was aligned against itself using NUCmer, implemented in the MUMmer suite v4.0.1^53^, with the --maxmatch option to identify all matches and the --nosimplify option to retain shadowed clusters. The minimum match length and minimum cluster length were set to 50 and 500 base pairs, respectively. Candidate ITRs were defined as inverted alignments connecting the two genome termini, with each aligned region occurring within 100 base pairs of the corresponding genome terminus, an alignment length of at least 1,000 base pairs, and nucleotide identity of at least 90%. The longest aligned regions were designated as the ITRs.

### Gene functional enrichment analysis

For protein domain gain and loss analysis, we used InterProScan v5.72-103.0^54^ to search for 23,794 Pfam protein domains, retrieved from the InterPro database release 103.0^55,56^, in the proteins encoded by all analyzed poxviruses. Protein domain gain and loss events for each ortholog group were then estimated similarly to ortholog group gain and loss events by evaluating the presence or absence of each protein domain in proteins encoded by orthologs in each ortholog group across all poxviruses. Functional enrichment analysis of protein domains was performed using Fisher’s Exact test followed by Benjamini–Hochberg correction.

### Statistical tests for pairwise comparisons and correlations

All statistical tests were performed in the R environment v4.5.1^57^ using the rstatix R package v0.7.3^58^. Normality and homogeneity of variances of data were assessed using the Shapiro–Wilk test and Levene’s test, respectively, with a significance threshold of 0.05 for both tests. Based on these test results, the Mann-Whitney U test was selected for pairwise comparisons of group medians. Correlation among parameters was quantified using Spearman’s rank correlation coefficient, whereas correlation among phylogenetically independent contrasts was assessed using the pic function implemented in the ape R package v5.8.1^59^.

## Data management and visualization

Data management and visualization were performed using the R environment. Figure plots were generated using the ggplot2 R package v4.0.0^60^. Phylogenetic trees were visualized using the ggtree R package v3.6.2^61^. Heatmaps were created using the ComplexHeatmap R package v.2.14.0^62^. Synteny plots were generated using the gggenomes R package v1.1.2^63^.

## Data and code availability

Computational codes used in this study are available in the GitHub repository (https://github.com/virus-informatics/Poxv_evol).

## Author Contributions

Project conception and supervision: Jumpei Ito

Data collection, generation, and analysis: Arnon Plianchaisuk

Figure preparation: Arnon Plianchaisuk

Support for evolutionary and phylogenetic analyses and result interpretation: Spyros Lytras

Support for bioinformatics analyses and result interpretation: Jumpei Ito, Spyros Lytras, Hiroyuki Hikida, Yuichiro Nakatani, Emma F. Harding, Aris Katzourakis, and Kei Sato

Manuscript drafting: Arnon Plianchaisuk

Manuscript editing: Jumpei Ito, Arnon Plianchaisuk, and Spyros Lytras

All authors reviewed and approved the final manuscript.

## Acknowledgments

The super-computing resource was provided by the Human Genome Center at The University of Tokyo. This research was supported by AMED SCARDA Center for Advanced Modalities and Drug Delivery Systems “CAMaD” (JP223fa627002, to Jumpei Ito); AMED SCARDA Japan Initiative for World-leading Vaccine Research and Development Centers “UTOPIA” (JP223fa627001, to Jumpei Ito, Kei Sato, Arnon Plianchaisuk, Spyros Lytras); AMED Japan Program for Infectious Diseases Research and Infrastructure (Interdisciplinary Cutting-edge Research) (JP26wm0325087, to Jumpei Ito); JST PRESTO (JPMJPR22R1, to Jumpei Ito); JSPS KAKENHI Grant-in-Aid for Scientific Research B (JP25K00116, to Jumpei Ito); AMED ASPIRE Program (25jf0126002, to Kei Sato); AMED SCARDA Program on R&D of new generation vaccine including new modality application (253fa727002, to Kei Sato); AMED Research Program on Emerging and Re-emerging Infectious Diseases (24fk0108907, 25fk0108690, to Kei Sato); AMED Japan Program for Infectious Diseases Research and Infrastructure (Collaborative Research via Overseas Research Centers) (25wm0225041, to Kei Sato); JSPS KAKENHI Grant-in-Aid for Scientific Research A (JP24H00607, to Kei Sato); the Platform Project for Supporting Drug Discovery and Life Science Research (Basis for Supporting Innovative Drug Discovery and Life Science Research (BINDS)) from AMED (JP24ama121012, supporting numbers S02820001 and S02820002, to Kei Sato). We also thank Dr. Amjad Khalaf, Prof. Elliot J. Lefkowitz, and Dr. Junna Kawasaki for constructive feedback on our manuscript development.

## Supplementary Tables

**Supplementary Table 1.** List of poxvirus genomes used in this study

**Supplementary Table 2.** Accession number of proteins from ortholog groups used in reconstructing a phylogenetic tree of poxvirus

**Supplementary Table 3.** Putative donors of HGT-derived ortholog groups

**Supplementary Table 4.** Numbers of gain and loss events of Pfam protein domains accompanied by ortholog group gain and loss

**Supplementary Table 5.** Enrichment analysis of protein domains from ortholog groups gained or lost during poxvirus evolution

**Supplementary Table 6.** Information on virus–host relationships and genomes of hosts or their closely related animals available in public databases used in this study

## Supplementary Figures

**Supplementary Figure 1.** Variation in gene number and genome size among poxviruses (A) Number of ortholog groups in each poxvirus genome. The dashed line represents the mean number of ortholog groups across all analyzed poxviruses. (B) Phylogenetically independent contrast analysis. The scatter plot shows the relationship between the contrasts in poxvirus gene number and those in genome size. *r*: Pearson’s correlation coefficient; *P*: *P* value of the correlation test; line: linear regression line. (C) Spearman’s rank correlation coefficients between virus and host genomic parameters, and Pearson’s correlation coefficients of phylogenetically independent contrasts for the same parameter pairs. No outliers were excluded. Asterisk: *P* value < 0.05. (D) Scatter plot showing the relationship between the number of poxvirus genes and the size of host genomes. Poxviruses associated with multiple host species were assigned the average genome size across their documented hosts. The fitted linear regression line and Spearman’s rank correlation coefficient (ρ) are shown. Melanoplus sanguinipes entomopoxvirus was excluded from the correlation and linear regression analyses due to the extremely large genome size of its host, desert locust (*Schistocerca gregaria*)^65^.

**Supplementary Figure 2.** Gene gain, duplication, and loss during poxvirus evolution (A and B) Numbers of ortholog group gain (A) and loss (B) events in each phylogenetic branch. Unlike Figure 2E, results are shown separately for each of the three ancestral state reconstruction methods. Branch colors indicate event counts on a pseudo-logarithmic scale. (C) Cumulative net changes in ortholog group number from the MRCA to each tip, estimated by each of the three ancestral state reconstruction methods.

**Supplementary Figure 3.** Genome architecture of poxviruses. (A) Central and terminal regions of poxvirus genomes. Central and terminal regions of each poxvirus genome that could be identified by the sliding window technique are shown in blue and pink, respectively. Orange, navy blue, and light grey arrows indicate gained, pre-existing, and undeterminable ortholog groups, respectively. Genomic positions are shown relative to the genome center, which was set to 0. (B) Duplicated genes in all ortholog groups in all poxvirus genomes. Genomic positions are shown relative to the genome center, which was set to 0. Line colors indicate genomic distribution of paralogous relationships between duplicated genes. Duplicated genes in ortholog groups of *N1R*/*p28* family and variola virus *B22R* are outlined in brown and magenta, respectively. Terminal genomic regions are shaded in pink. ITRs are shown by yellow ribbons.

**Supplementary Figure 4.** Host–donor taxonomic associations of genes horizontally transferred to poxvirus. For each virus, the proportion of ortholog groups assigned to each putative donor taxon was calculated relative to the total number of ortholog groups in that virus. Heatmap colors represent these proportions. The taxon of the documented host of each poxvirus is indicated by the letter H. Only the results for the top three taxa of putative donors are shown for each poxvirus.

**Supplementary Figure 5.** Number of protein domain loss events inferred for each branch of the poxvirus species tree. Only those accompanied by ortholog group gain events were counted. Branch colors represent event counts on a pseudo-logarithmic scale.

**Supplementary Figure 6.** Number of poxvirus proteins that could be aligned with non-poxvirus proteins, stratified by percentage query coverage per high-score segment pair. Bin size: 10; dashed line: median.

