## Supplementary figures and images for "Poxvirus gene repertoires are shaped by expansion-biased birth-and-death dynamics and genome architectural constraints"

### Supplementary Figure 1

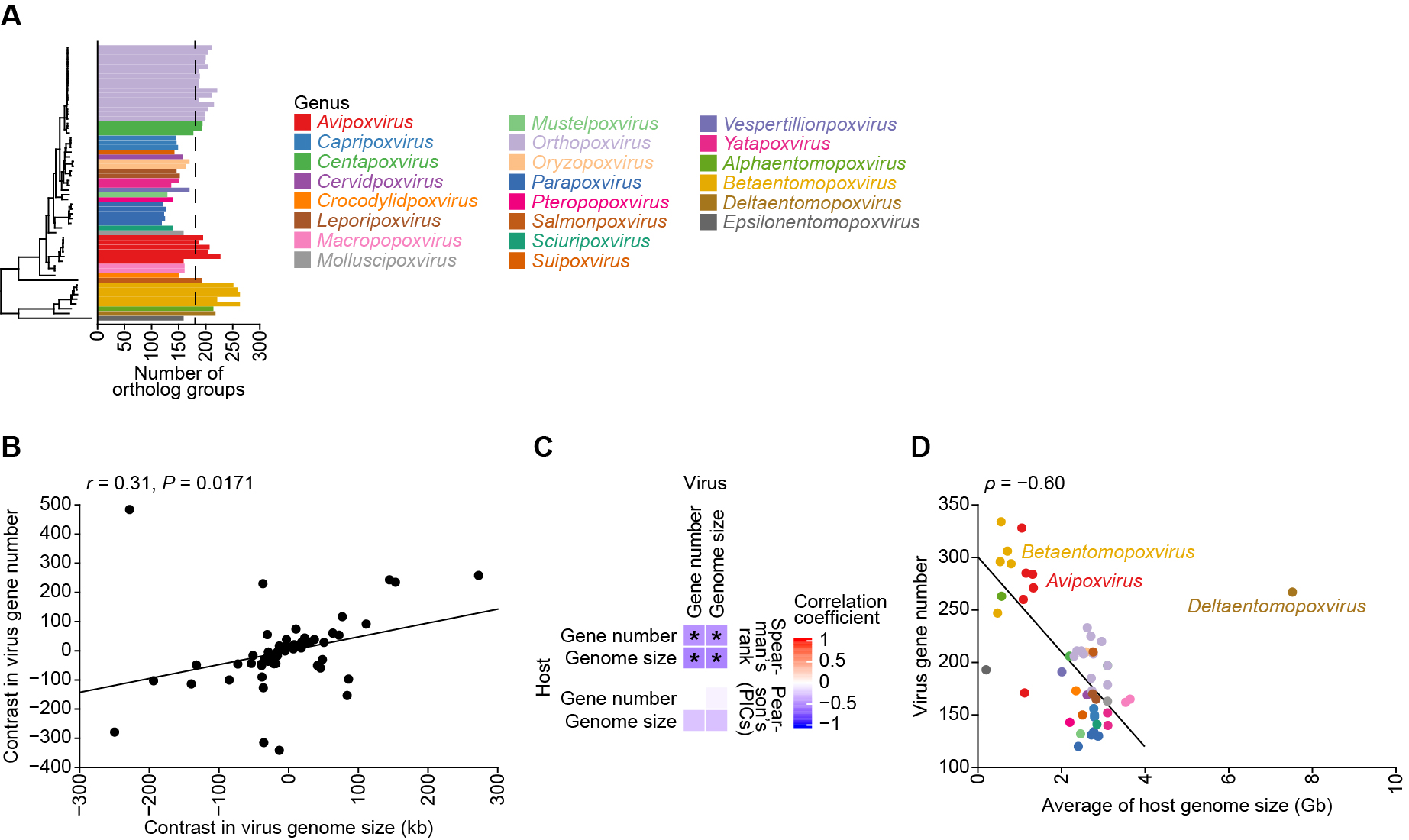

### Supplementary Figure 2

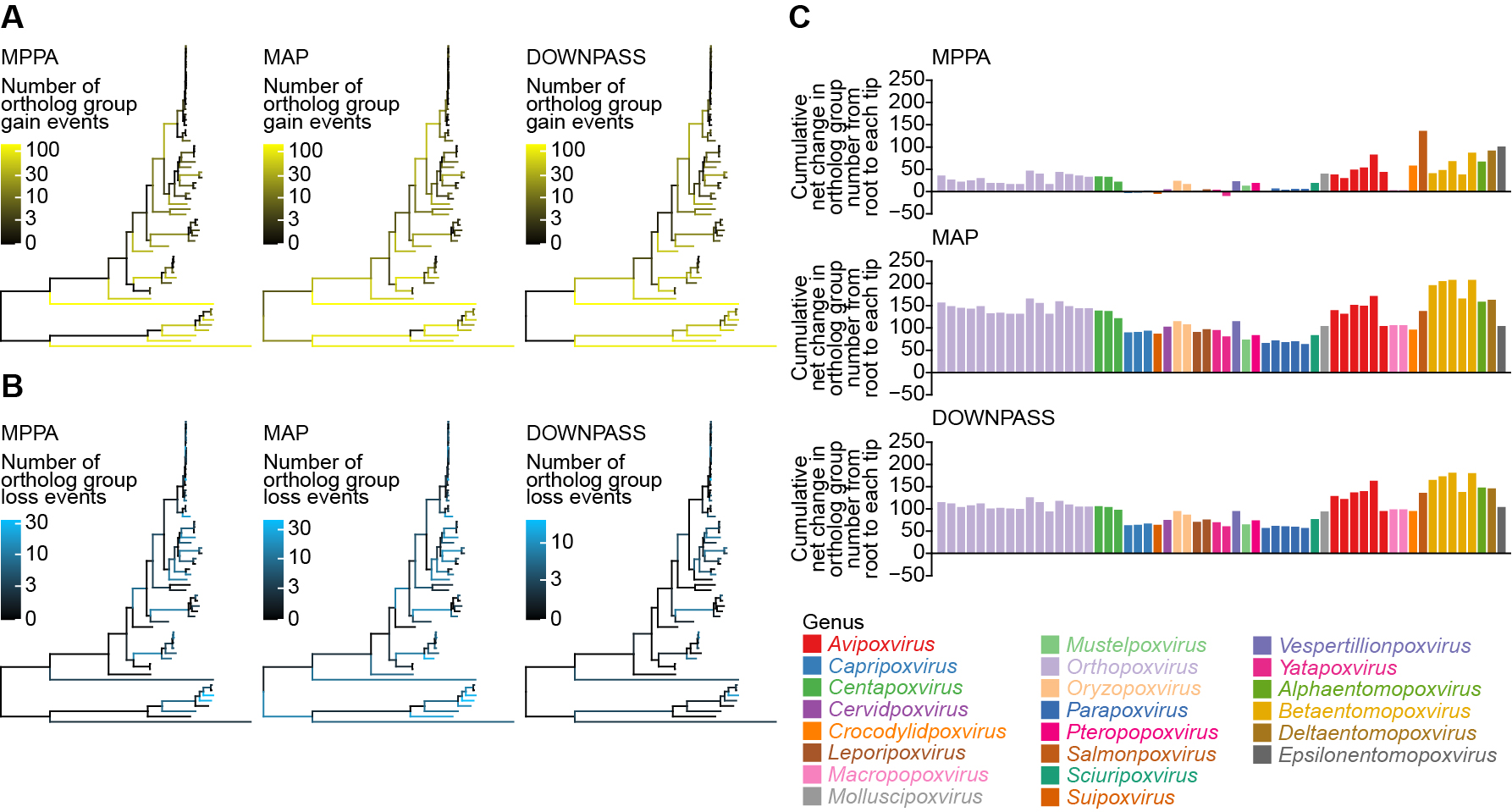

### Supplementary Figure 3

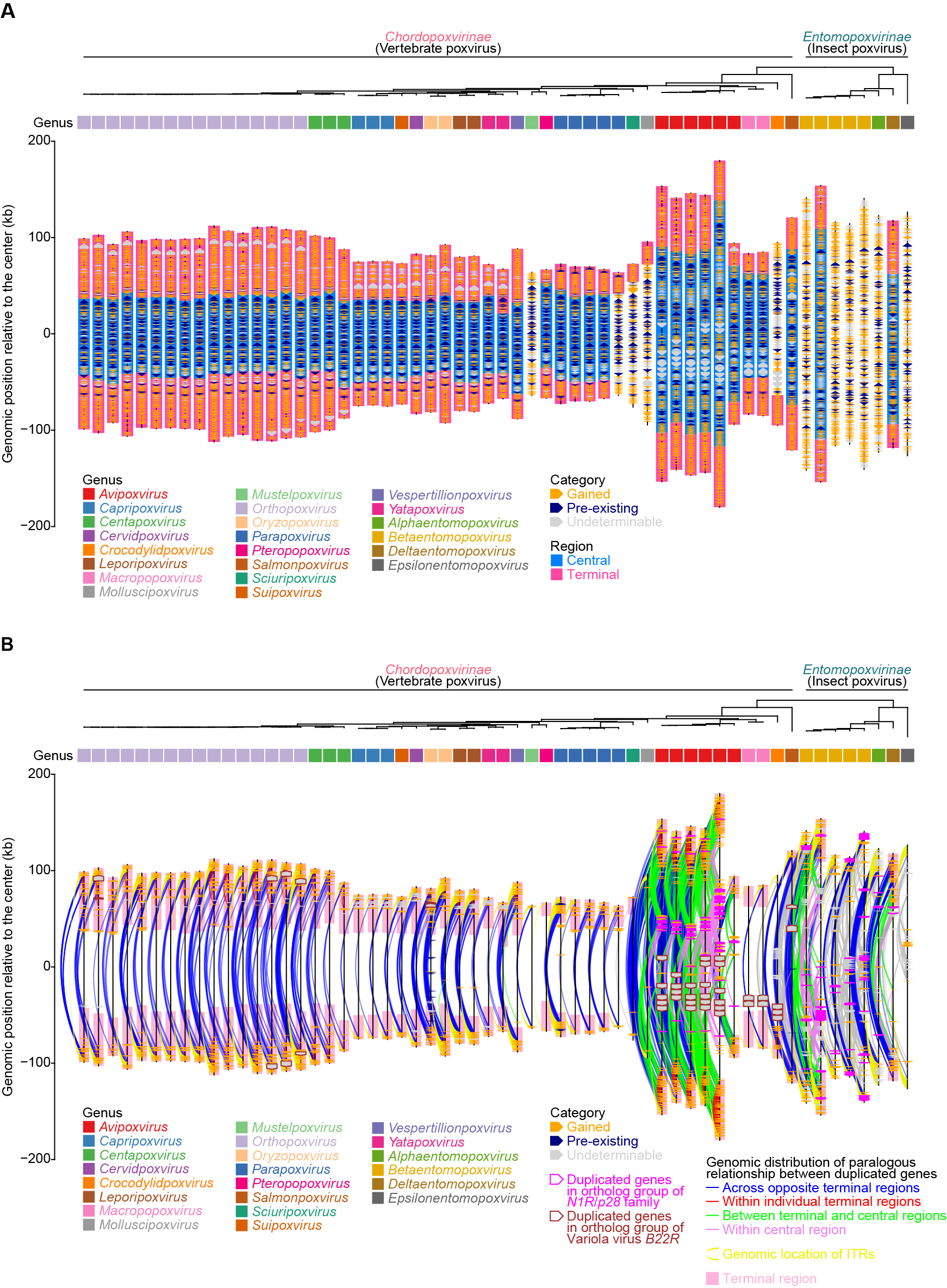

### Supplementary Figure 4

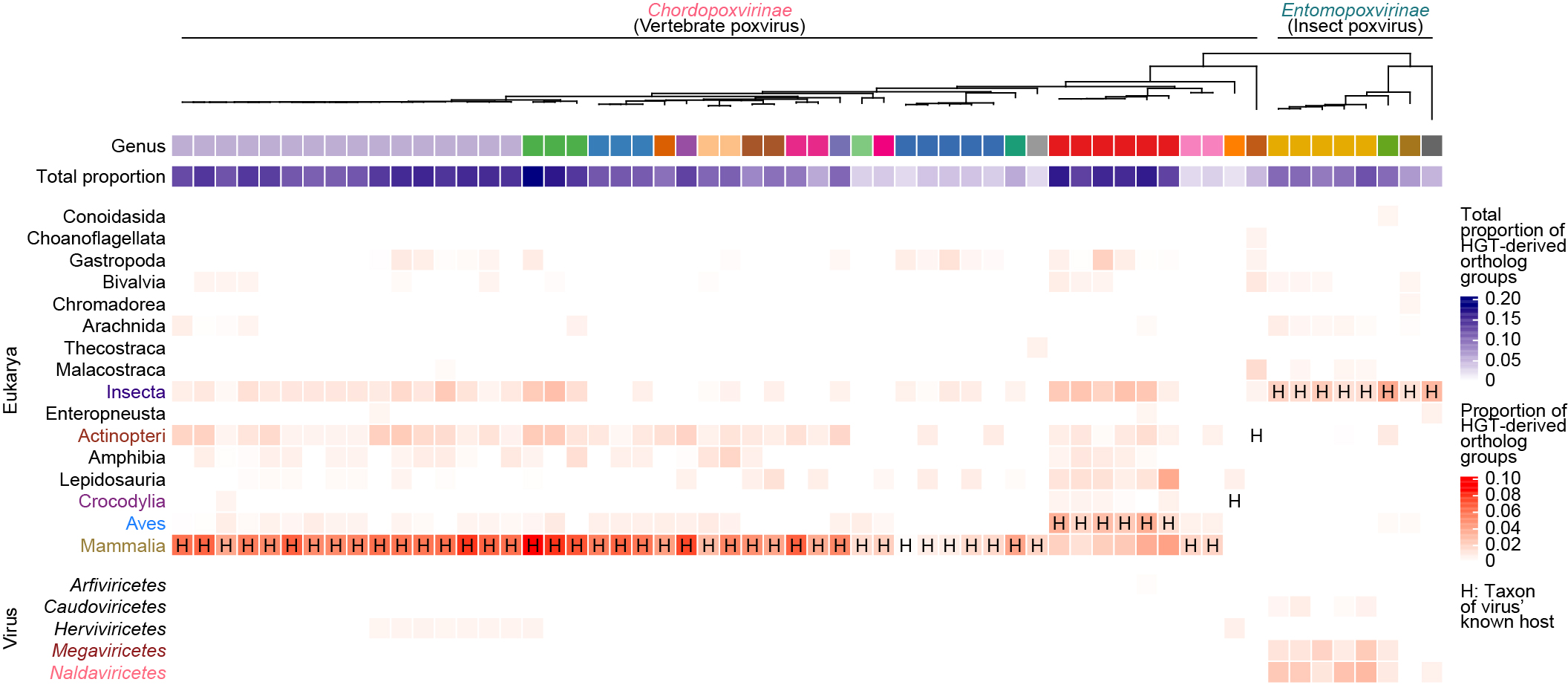

### Supplementary Figure 5

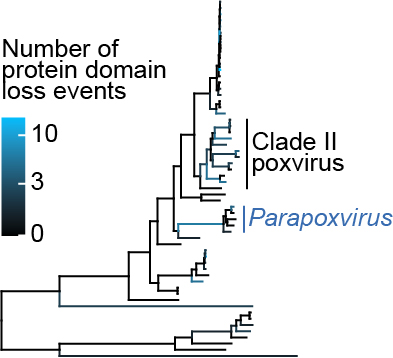

### Supplementary Figure 6

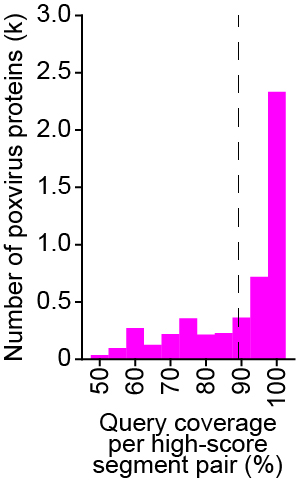
